# Targeted memory reactivation in human REM sleep elicits detectable reactivation

**DOI:** 10.1101/2021.12.01.470530

**Authors:** Mahmoud E. A. Abdellahi, Anne C. M. Koopman, Matthias S. Treder, Penelope A. Lewis

## Abstract

Several studies show that memories are reactivated during non-rapid eye movement (NREM) sleep, but the question of whether equivalent reactivation can be detected in rapid eye movement (REM) sleep is hotly debated. To examine this, we used a technique called targeted memory reactivation (TMR) in which sounds are paired with learned material in wake, and then re-presented in subsequent sleep to trigger reactivation. We then used machine learning classifiers to identify reactivation in REM related to the encoded wake activity. The reactivation we measured was mediated by high theta activity and was sometimes temporally compressed and sometimes dilated compared to wakeful experience. Reactivation strength positively predicted overnight performance improvement. These findings provide the first evidence for wake-like memory reactivation in human REM sleep after TMR.

## Introduction

The reactivation of memories in non-REM sleep is widely supported by evidence from humans, rodents, and other animals(1–4), but it is still unclear whether equivalent reactivation occurs in REM sleep. The first evidence for non-REM reactivation came from rodents(5,6). In humans, NREM reactivation has been identified using EEG classifiers(3,7–9), fMRI(10,11), and intracranial recording(12). Turning to REM sleep, only a few rodent studies have shown evidence for reactivation(5,13–15). In humans, EEGs of REM sleep occurring after training on two different tasks have been elegantly distinguished (16) but, the reinstatement of wake-like memory patterns in REM sleep has yet to be demonstrated.

Targeted memory reactivation (TMR), a technique which allows the active triggering of memory reactivation, is linked to both neural(17) and behavioural(18–20) plasticity when applied in non-REM sleep. TMR provides a set of temporal windows when reactivation is biased by an external cue, and this can be used to evaluate the detectability of reactivation. In the current study, we use TMR to build on the work of Schönauer and colleagues (2017) by looking for wake-like task related activity after TMR cues during REM. We thus search for a direct link between brain activity associated with a memory in wake and cued reactivation in REM.

Theta activity predominates during REM sleep(21,22), and human studies suggest a relationship between memory and theta(23,24). Synchronisation of theta has been shown to predict encoding ability(25), and wakeful theta is suggested to provide a preferable window for the encoding of new information(26–28). Event-related changes in spectral theta power during encoding have also been shown to tag memories for selective consolidation(29). Interestingly, two of the rodent studies that have supported memory reactivation in REM, showed a link between such reactivation and theta activity(13,15). We were therefore interested to determine whether theta is associated with TMR cued reactivation in human REM sleep.

We chose the serial reaction time task (SRTT), which is known to be sleep sensitive(30,31), to examine these questions. The SRTT is modulated by TMR in NREM sleep(32–34) has also been strongly linked to REM(35–37). For instance, brain areas which were activated during the execution of a serial reaction time task have been shown to reactivate during subsequent REM(35). Results from another serial reaction time study suggest that regional cerebral reactivation in REM sleep reflects the reprocessing of learned material(36). Furthermore, greater connectivity between premotor cortex, posterior parietal cortex, and bilateral pre-supplementary motor has been observed during REM after training on this task(37). These studies suggest that reprocessing of learned material is taking place in REM. However, direct evidence linking wake and sleep patterns is still missing.

In our SRTT, participants were presented with audio-visual cues and responded by pressing four buttons (two from each hand). Cues were organised into a twelve-item sequence. Sounds were replayed during subsequent REM to trigger the associated memories of left- and right-hand presses (Fig. 1). We used two sequences and replayed only one of them in sleep. For control, we also included an adaptation night in which participants slept in the lab and the same tones that would later be played during the experimental night were played. This provided data in which tones could not have evoked memory reactivation, as participants had not yet learned the behavioural task, see methods for more details about the SRTT. We used these data to discriminate between neural reactivation of left- and right-hand button presses using linear classification on EEGs during REM sleep. After detecting reactivation, we examined key characteristics of this TMR cued reactivation, including association with theta activity, temporal compression/dilation.

**Fig. 1.**
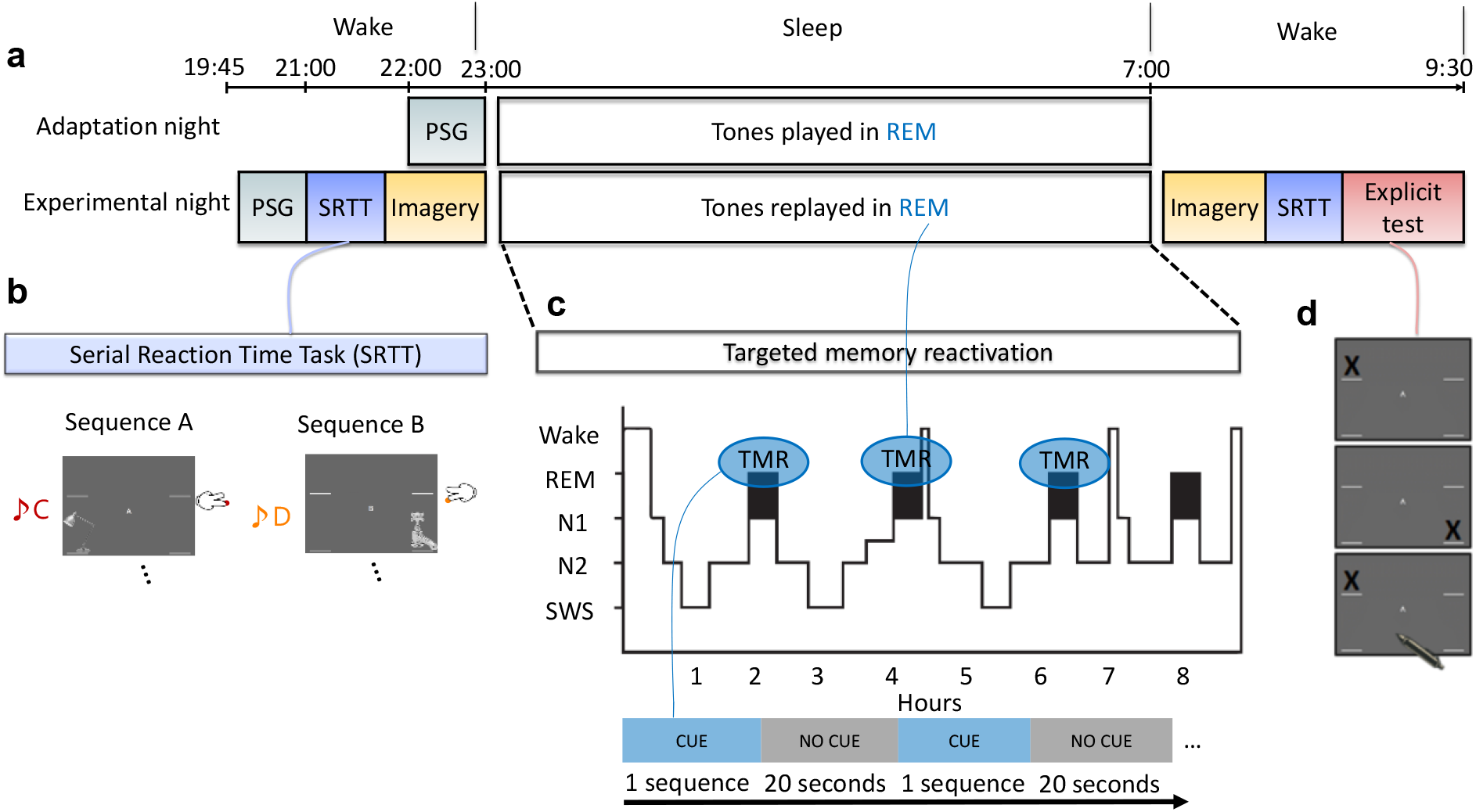
Experimental design. **a** The experiment consisted of two nights an adaptation and an experimental night. In the adaptation night, tones were presented to the participants during REM sleep and EEG recordings were acquired. In the experimental night, participants were wired-up for EEG then completed the serial reaction time task (SRTT) and an imagery task in which they were cued with pictures and sounds, but only imagine performing the finger tapping (without movement). Afterwards, participants slept in the lab and TMR cues were presented. After waking up, participants completed the motor imagery and SRTT again, and finally they did the explicit recall task. **b** In the SRTT, images are presented in two different sequences each with a different set of tones. Each image is associated with a unique tone and requires a specific button press. In the imagery task, participants hear the tones and see the images as in the SRTT, but only imagine pressing the buttons. This imagery data was used for classification as it has cleaner signals compared to SRTT since there are no movement artifacts. **c** The sounds of only one learned sequence (cued sequence) were played during REM sleep, with a 20 second pause between repetitions. **d** Participants were asked to mark the order of each sequence on paper as accurately as they can remember during the explicit recall test after sleep.

## Results

### Detection of memory reactivation after TMR cues

We trained our classification models using sleep data and then tested them on wake data. Training a classification model on wake could cause the model to weigh features which are dominant in wake very highly even if those features were absent from sleep. By training classification models on sleep data, we ensured that the features associated with reactivation were used by the models, and the models were thus able to look for these in the stronger, less noisy, signals recorded during wake. Because we had only 16 electrodes, we used a searchlight approach to locate the channels of interest, see methods for details. This showed that classification was highest near the motor area (Supplementary Fig. 5).

We used a linear discriminant analysis (LDA) classifier in a time x time classification procedure(38), see methods for details. We repeated the classification process using the adaptation night to be certain that the classification was not caused by sound induced effect or EEG noise rather than reactivation of the encoded memory. We compared the results from the two nights, both to each other and to chance level using cluster-based permutation. In the adaptation night, no significant classification strength was found compared to chance (area under the receiver operating characteristic curve (AUC) = 0.5), demonstrating that classification of this control condition did not differ from chance level. By contrast, comparison of the experimental night against chance showed a significantly higher AUC for the experimental night (Fig.2a) which occurred around 1 sec. after the onset of the cue at time 0 (Fig. 2b). Comparison of the experimental night to the adaptation night also showed a significantly higher AUC for the experimental night, described by a cluster in the same timeframe (Fig. 2a). This clearly demonstrates that we can detect memory reactivation after TMR cues in REM sleep, and that we can discriminate between right- and left-hand movements during REM sleep. These findings are significant when evaluated against both control and chance level.

**Fig. 2.**
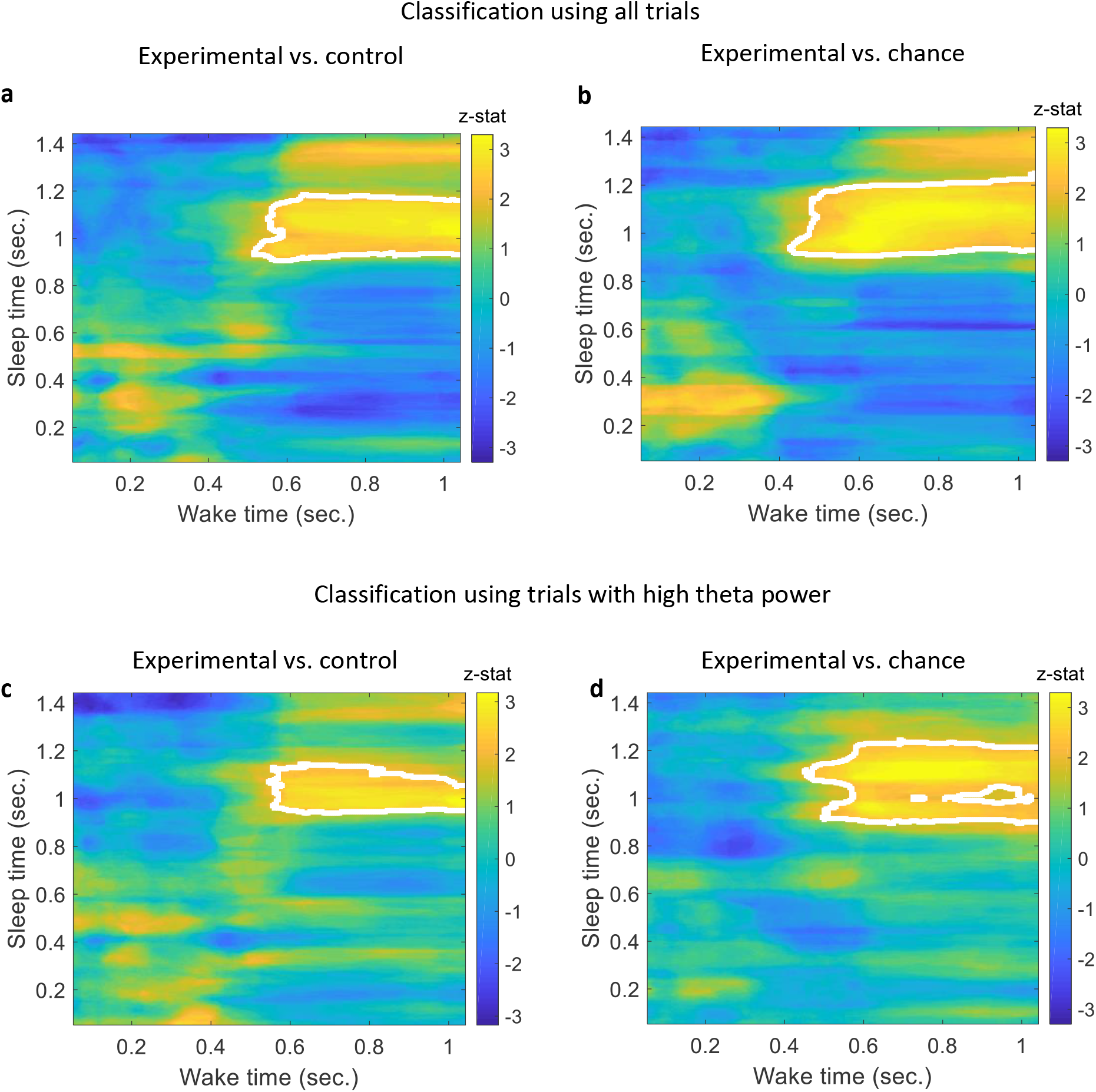
Classification of left-vs. right-hand. **a** A comparison between the classification performance of the experimental vs. adaptation night using all trials reveals a significant effect described by a cluster which shows a higher classification performance for the experimental night compared to adaptation (n= 14, p= 0.01), a z-statistic value at every point is shown and cluster edges are marked with white after correcting for multiple comparisons with cluster-based permutation (see methods for details). **b** Classification performance for the experimental night was also significantly higher than chance (AUC: 0.5) as shown by the cluster after correction (n= 14, p< 0.0001). **c** A comparison between the experimental and adaptation night classification using trials with high theta power reveals a significant effect described by a cluster that shows a higher classification performance for the experimental night compared to adaptation (n= 14, p= 0.028). **d** A comparison between the classification of experimental night using trials with high theta power and chance level shows a significant effect described by the cluster after correction (n= 14, p= 0.001).

### High theta activity mediates reactivation

To test for a relationship between theta activity and reactivation we performed a median split on theta power, creating two groups of trials for each participant: those with high theta power and those with low theta power, see methods for details. This split was performed for both experimental and adaptation nights. We then compared the classification results of each half of the median split (high and low theta power trials) in the experimental night both to chance level and to the equivalent high or low theta power trials in the adaptation night. For high theta trials, this showed a significant effect (Fig. 2c), explained by a cluster occurring around the same time as in the classification result using all trials, (Fig.2a-b), (n = 14, p = 0.028), there was also a significant difference from chance level (Fig. 2d), (n = 14, p = 0.001). A comparison between high and low theta trials within the experimental night showed no significant difference, this could be due to the high variance in low theta trials which is seen when we compare classification of low theta trials to chance level and adaptation night which showed no significant effect (Supplementary Fig. 4). These findings demonstrate an association between high theta activity and reactivation.

To determine whether theta activity was the actual feature allowing the discrimination of classes, we band-pass filtered the signals in the theta band and re-ran our classification analysis. Interestingly, classification did not differ from chance in this filtered data (p > 0.4) suggesting that while high theta activity offers a preferred window for reactivation, theta activity itself is not the information used to detect reactivation. Time x time classification AUC plots using all trials and also using high theta power trials are shown in (Supplementary Fig. 3).

Prior work from our group has suggested that TMR’s ability to elicit classifiable NREM reactivation in the SRTT task declines across the stimulation period(7). To examine this in our current dataset, we analysed whether trials with high theta activity came from anytime of stimulation or were specific to the beginning or the end of stimulation period. We indexed stimulations to have the range 0 to 1 with 0 the first stimulation and 1 the last one. This showed that trials with high theta activity were not specific to the beginning or end of the stimulation period when compared to the middle of stimulation time 0.5 (Wilcoxon signed rank test, n = 14, p = 0.55). To be sure that the recording quality and any other noise were not causing the classification seen with high theta activity, we performed classification after filtering in different frequency bands. In this control, we used three different bands, lower and higher than theta and a broad range of frequencies: [0.5 3] HZ, [9 16] HZ, and [0.5 30] HZ. The signals were band pass filtered in these bands and we performed the same median split as performed on theta band, but none of the classification using these bands produced significant cluster(s). This clearly demonstrates that theta activity mediates reactivation and this is unique to theta band.

### Correlation of classification performance with behaviour

We next tested for a relationship between classification performance and improvement on the behavioural task using a spearman correlation, see methods for details. This revealed a positive correlation (Fig. 3a; n = 14, r = 0.74, uncorrected p = 0.0035 and Bonferroni corrected p = 0.01). This indicates that more strongly detected reactivation was associated with greater overnight sequence improvement. Importantly, this correlation only existed for the reactivated sequence (for non-reactivated: n = 14, r = 0.24, uncorrected p = 0.4). This finding suggests that the degree of detecting reactivation in REM positively predicts the extent to which sequence memory improves overnight. Further details on behaviour can be found in the discussion.

**Fig. 3.**
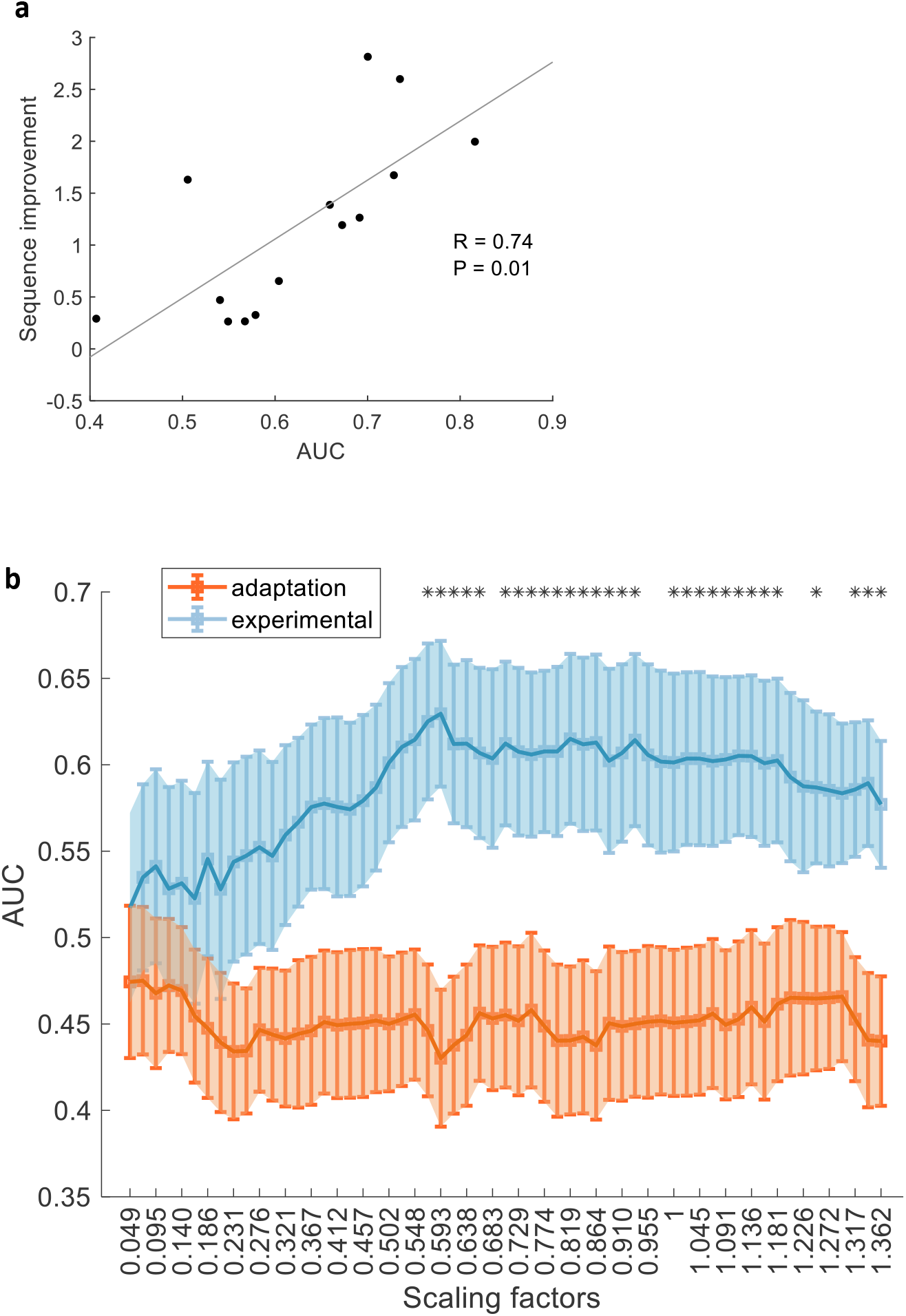
Characteristics of detected reactivation. **a** Classification performance positively correlated with behavioural improvement for the reactivated sequence (spearman correlation, n= 14, r= 0.74, Bonferroni corrected p= 0.01). **b** Classification results for different scaling factors (SF), orange curve for the adaptation night and blue for the experimental night, significant difference (p<0.05) is marked with an asterisk. This shows that reactivation could be compressed (~0.6 times wake activation) or dilated (~1.4 times wake activation).

### Temporal compression/dilation of reactivation

Prior work has shown that reactivation in non-REM sleep is often temporally compressed with respect to wake(4,39,40). A recent finding showed that wake and NREM1 reactivation could be sometimes compressed and sometimes dilated within the same data(41). This motivated us to determine whether reactivation in REM occurs for the same amount of time as the original experience in wake. Thus, we performed an analysis of temporal compression/dilation. We tackled this problem within a machine learning context such that we were able to compare the classification results to chance level and the control condition. We thus looked for reactivation strength at different scaling factors by varying the window length we extract from REM sleep. The scaling factor represents the length of the small window from sleep divided by the length of wake trial. We take a small window from sleep time (e.g., 10 time points) and then resize it to match the length of wake trial (1.1 sec.), see methods for more details. A classifier is then trained on the resized sleep windows and tested on wake and a classification performance is obtained. This is repeated for every participant at each scaling factor, Fig. 3b.

## Discussion

We have demonstrated that reactivation of wake-like neural activity after TMR cues can be detected in human REM sleep using EEG classifiers. Such reactivation appears to be most prominent about one second after the sound cue onset. Reactivation is associated with high theta power, which appears to provide a preferred window for this activity, although it does not carry the discriminative information needed to actually detect the reactivation. Interestingly, reactivations can be compressed or dilated compared to the encoded pattern in wake.

### Temporal structure of reactivation

Memories have a different temporal structure during reactivation compared to encoding. For instance, in rodents, replay has been shown to occur at a faster rate than in the original task in both wake and SWS(39,42,43). However, the exact differences in timing are unclear, since some studies of rodent non-REM sleep showed a compression of 6 to 7 times compared to wake(40), while other studies showed compression rates varying between 10 and 20 times compared to wake(4,39). Meanwhile, REM replay in rodents appears to be almost the same length as wakeful activity, or even to be dilated with respect to wake (13). Turning to humans, wake and NREM1 reactivations are sometimes compressed and sometimes dilated(41). Our analyses are in keeping with this, since we show that reactivation occurs in varying lengths and could sometimes appear compressed: almost twice as fast as wake activation and sometimes dilated: ~1.4 the length of wake activation, Fig. 3b.

Detectable reactivation is temporally consistent among trials and participants, this is reflected in the timing of the clusters found in Fig. 2. During wake, classification is more prominent around 0.6 seconds after cue, whereas in REM it starts about 0.4 seconds later than that. This delayed start could potentially be due to the brain taking more time to process information and reactivate a memory during REM sleep than during wake. In fact, reactivation in SWS has also been shown to be delayed, appearing about 1 second after cue onset(3) and sometimes later around 2 seconds from cue onset(9).

### Theta activity

REM sleep is dominated by theta activity, which has been linked to reactivation(13) and is thought to support the consolidation process(44). Theta is linked to attention during wake(45) and is more prominent with higher executive control(46). Wakeful theta is also associated with the encoding of new information and memory processing(28). Furthermore, neuronal firing relative to theta phase has been shown to impact upon whether synapses are strengthened or weakened, since stimulation of the positive theta phase induces long-term potentiation and stimulation of the negative theta phase induces depotentiation(47). Increased theta power has previously been shown to be important for successful cueing during sleep(48), and theta phase consistency indicated reactivation after TMR cues(8). In our data, trials with higher theta power show greater evidence of memory reactivation. Theta therefore appears to be a marker of reactivation but does not embody the reactivation in and of itself. It could, for instance, provide a timing function for reactivation.

### Correlation with behaviour

The performance of our REM classifiers positively predicted overnight improvement on sequence memory. Similar correlations between classification performance and behaviour have been found in non-REM TMR(3,10,12). This could lead to the speculation that more reactivation means more consolidation, and therefore better post-sleep performance. However, TMR in REM sleep did not lead to overall benefit in performance either in our current dataset(49) or in previous studies(11,50). Furthermore, examination of natural (un-cued) sleep showed a correlation between overnight memory improvement and the extent of memory-linked activity identified in NREM but not REM sleep(16) potentially suggesting a difference in the function of replay in these two sleep stages. This absence of behavioural improvement could be due to individual differences in the way REM processes this task, potentially relating to an interaction between the extent or style of learning and REM processes. The benefit of cueing could also appear after a longer delay; see (34) for delayed impacts of TMR in SWS on this task.

### Parallel characteristics of reactivations in REM and NREM

While there is already a growing body of literature about TMR cued reactivation in non-REM sleep(1–3,9,51), our findings provide initial information about human memory reactivation using TMR in REM. In fact, our data suggest several parallels between reactivation in these two sleep stages. For instance, similar to non-REM(9), TMR-locked reactivation in REM is delayed compared to wake after cue onset. Furthermore, reactivation in REM is related to the oscillatory structure of sleep (e.g., theta activity), paralleling the relationship between reactivation in non-REM and graphoelements like slow oscillations(52), spindles(9,53), and ripples(12). Importantly, the reactivations we detected appeared to occur in roughly the same area of the brain as is used in performing the task, and the extracted features of reactivation are similar to those extracted during wake activation which is why it is detectable using our machine learning models. Notably, REM reactivations differ temporally from the encoded pattern, and appear to be sometimes compressed and sometimes dilated. This parallels reactivation in NREM1 that was both compressed and dilated(41).

## Conclusion

The question of whether memories reactivate in REM as well as non-REM sleep has been debated for some years. REM reactivation has been suggested by modelling(54) and evidence of learning dependent activation in human REM sleep has been observed (35,36). However, null behavioural findings from human REM TMR studies(1,11) have led to scepticism amongst sleep researchers. Our current findings provide clear evidence that TMR in REM elicits detectable reactivation. Furthermore, our analysis uncovers several important properties of REM reactivation, showing strong parallels with reactivation in non-REM sleep. Moreover, we see evidence of both compression and dilation which shows that reactivation temporally varies from wakeful experience. More work is needed to explore how such reactivation links to behavioural and neural plasticity, and how this differs across a variety of cognitive tasks.

## Methods

### Participants

EEG and behavioural data were collected from human participants (n = 16) (8 females, 8 males, and age mean: 23.6). Two participants were excluded due to technical problems (n = 14). All participants had normal or corrected-to-normal vision, normal hearing, and no history of physical, psychological, neurological, or sleep disorders. Responses in a pre-screening questionnaire reported no stressful events and no travel before commencing the study. All participants reported non-familiarity with the SRTT and all of them were right-handed. Participants did not consume alcohol in the 24 hours before the study and caffeine in the 12 hours prior to the study or perform any extreme physical exercise or nap.

### Experimental Design

The SRTT was adapted from(32) and participants performed SRTT before and after sleep. In SRTT, sounds cued four different finger presses. During wakeful encoding, participants learned two 12-item sequences, A and B, A: 1 2 1 4 2 3 4 1 3 2 4 3 and B: 2 4 3 2 3 1 4 2 3 1 4 1. The location indicated which key on the keyboard to press as quickly and accurately as possible afterwards the next image appeared, locations and buttons associations were: 1 – top left corner = left shift key; 2 – bottom left corner = left Ctrl; 3 – top right corner = up arrow; 4 – bottom right corner = down arrow. Sequences had been matched for learning difficulty; both contained each item three times. The blocks were interleaved so that a block of the same sequence was presented no more than twice in a row, and each block contained three repetitions of a sequence. There were 24 blocks of each sequence (48 blocks in total), after each block a 15-second pause which could be extended by participants if they wish, during the pause participants were informed with their reaction time and error rate for the last block. A sequence was paired with a group of pure musical tones, either low tones within the 4th octave (C/D/E/F) or high tones within the 5th octave (A/B/C#/D) (counterbalanced). The 48 blocks of sequences A and B were followed by four blocks that contained random sequences. Two of these blocks were paired with the tone group of one sequence that was later replayed in REM sleep, and the other two were paired with the tone group of the other non-reactivated sequence. On the centre of the screen ‘A’ and ‘B’ appeared while participants performed the task, and they knew that there were two sequences. However, they were not asked to explicitly learn the sequences. The first sequence to present as well as the sequence to reactivate was counterbalanced across participants. At the beginning of each trial at time 0, a 200 ms tone was played, and at the same time a visual cue appeared in one of the corners of the screen. Participants were instructed to keep individual fingers of their left and right hand on the left and right response keys, respectively. Visual cues were neutral objects or faces, (Supplementary Fig. 1), used in previous study(32). Visual cues appeared in the same position for each sequence (1 = male face, 2 = lamp, 3 = female face, 4 = water tap). Visual cues stayed on the screen until the correct key was pressed, after which an 880 ms inter-trial interval followed. After completing the SRTT, participants performed the imagery task by only imagining pressing the buttons, without movement. This task consisted of 30 interleaved blocks (15 of each sequence) and was presented in the same order as during the SRTT. This imagery data is used for classification as it has higher signal to noise ratio since there is no movement compared to actual finger presses. Each trial consisted of a tone and a visual stimulus, the latter being shown for 270 ms and followed by an 880 ms inter-trial interval. There were no random blocks during the imagery task and no performance feedback during the pause between blocks. As a control, participants were asked to sleep in the lab before doing the SRTT training. The control night followed the same criteria as the actual experiment with the difference that the sounds were not associated with any task. Participants performed the task again in the morning but this time motor imagery then SRTT. They were then asked to try to remember the locations of the images of the two sequences to test their recall for each of the sequences.

### EEG recording and pre-processing

EEG signals were acquired from 21 electrodes, following the 10-20 system. Thirteen electrodes were placed on standard locations: FZ, CZ, PZ, F3, F4, C5, CP3, C6, CP4, P7, P8, O1, and O2. Other electrodes were: left and right EOG, three EMG electrodes on the chin, and the ground electrode was placed on the forehead. Electrodes were referenced to the mean of the left and right mastoid electrodes. The impedance was <5 kΩ for each scalp electrode, and <10 kΩ for each face electrode. Recordings were made with an Embla N7000 amplifier and RemLogic 1.1 PSG Software (Natus Medical Incorporated). PSG recordings were scored by trained sleep scorers and only the segments scored as REM were kept for further analyses.

Data were collected at 200 Hz sampling rate. EEG signals were band pass filtered in the frequency range from (0.1 to 50 HZ). Subsequently, trials were cleaned based on statistical measures consisting of variance and mean. Trials were segmented −0.5 sec. to 3 sec. relative to the onset of the cue. Trials falling two standard deviations higher than the mean were considered outliers and rejected if they show to be outliers for more than 25% of the channels. If trials were bad in less than 25% of the channels, they were interpolated using triangulation of neighbouring channels. Thus, 9.8% and 10.5% of trials were considered outliers and removed from the experimental night data and the adaptation night, respectively.

Data was subsequently analysed with independent component analysis (ICA), to remove eye movement artifacts which can occur during REM. Components identified by the ICA that were significantly correlated with the signals from the eye electrodes (corrected for multiple comparisons) were removed. All channels for each participant were manually inspected and artifacts were removed. Because TMR will not be effective with all trials, we also rejected trials with low variance that do not differ from their mean across time since they are unlikely to contain a response. The number of clean trials kept after cleaning was consistent among participants such that they contribute equally to the group-level analysis and that number was 366 trials, it was determined according to the participant with the lowest number of such trials. All cleaning was done on all trials irrespective of cue information and stimulation night to avoid bias.

### Time x time classification with time domain features

We adopted a time x time classification approach after smoothing the EEG signals of motor imagery using 100 ms window such that every time point is replaced with the average of the 50 ms of both sides around it. A region of interest based on searchlight analysis was used to select channels of interest, this was done on different participants who performed the same task to avoid circularity. In the time x time classification, every time point from sleep was used to train LDA classifier, which was applied to all time points from wake in order to get one row of classification results in the time x time classification plot (e.g., Fig. 2a). The process was repeated for all time points after a sound cue in sleep (trial length in sleep was: 1.5 sec. and 1.1 sec. in wake) (Supplementary Fig. 2). We use the area under the receiver operating characteristic curve (AUC) as the performance measure in our classification. Analyses were done using FieldTrip(55), MVPA-Light toolbox in Matlab(56), and customised scripts using Matlab 2018a. The clustering window used for cluster-based permutation was the whole length of trials (i.e., the whole time x time classification duration) which gave a stringent test. In other words, we did not limit the clustering window to a specific time window.

### Theta power calculation

We calculated theta power using band pass filtering in the range (4 to 8 Hz) and Hilbert transform to extract the instantaneous power information of the signals. The power of a trial is calculated as the average power across all time points of that trial and all channels. We then divided the trials of every participant into two groups based on the median power of trials. This gave us trials with high theta power (higher than median) and low theta power (lower than median).

### Temporal compression/dilation of reactivation

We analysed the temporal compression and dilation by looking at reactivation strength at different scaling factors by varying the length of a sliding window on sleep. A scaling factor is calculated as the length of the small window from sleep divided by the length of wake trial, thus, a value smaller than 1 indicates compression in sleep. A window from sleep is resized to match the length of wake trial (1.1 sec.). We then use principal component analysis (PCA) to reduce the dimensionality of features and keep the PCs with the highest variance. We kept components containing 95% of the explained variance. All of the windows of a specific scaling factor are then used to train a LDA classifier and the classifier is then applied to wake and the classification AUC value is calculated. Thus, for each scaling factor a classification performance for each participant is obtained. Subsequently, scaling factors are compared to chance level of 0.5 as well as to classification from the control night.

### Correlation of classification performance with behaviour

Classification performance was averaged inside the cluster. AUC values from the high theta power classification were tested for correlation with the behavioural improvement. The behavioural improvement was calculated as: [(random blocks after sleep - the best 4 blocks after sleep) – (random blocks before sleep – the best 4 blocks before sleep)] / (random blocks before sleep – the best 4 blocks before sleep). The result was corrected for other measures using Bonferroni correction. We extracted three behavioural measures: early blocks improvement, late blocks improvement, best blocks improvement (described above). Early blocks improvement was defined as: [(random blocks after sleep - the first 4 blocks after sleep) – (random blocks before sleep – the last 4 blocks before sleep)] / (random blocks before sleep – the last 4 blocks before sleep). Late blocks improvement was defined as: [(random blocks after sleep - the last 4 blocks after sleep) – (random blocks before sleep – the last 4 blocks before sleep)] / (random blocks before sleep – the last 4 blocks before sleep).

### Correcting for multiple comparisons

Multiple comparisons correction was done using MVPA-Light toolbox in Matlab(56) and customised scripts. Cluster-based permutation testing was used with sample-specific threshold of 0.05. Permutation test threshold for clusters was 0.05, and 10,000 permutations were calculated.

## Data availability

All relevant data generated or analysed are available along with Matlab scripts. Data are available at the Open Science Framework (OSF):

https://osf.io/wmyae/?view_only=5bd3badf3acb46a88a209dbed57c1a85

https://osf.io/fq7v5/?viewonly=02380297e8334391ab9b473e4efe7d0c

## Ethics Statement

This study was approved by the School of Psychology, Cardiff University Research Ethics Committee, and all participants gave written informed consents. Information of the participants are anonymised.

## Acknowledgements

This work was supported by the ERC grant SolutionSleep to P.A.L and ERC funded the Ph.D. of M.E.A.A. We would like to thank Martyna Rakowska and Lorena Santamaria and all members of our group NaPs for the useful comments.

## Author contributions

A.C.M.K. and P.A.L. designed the experiment. A.C.M.K collected the data. M.E.A.A analysed the data.

M.E.A.A., P.A.L., A.C.M.K. and M.S.T wrote the manuscript.

