## Supplementary figures for "Targeted memory reactivation in human REM sleep elicits detectable reactivation"

### Supplementary material

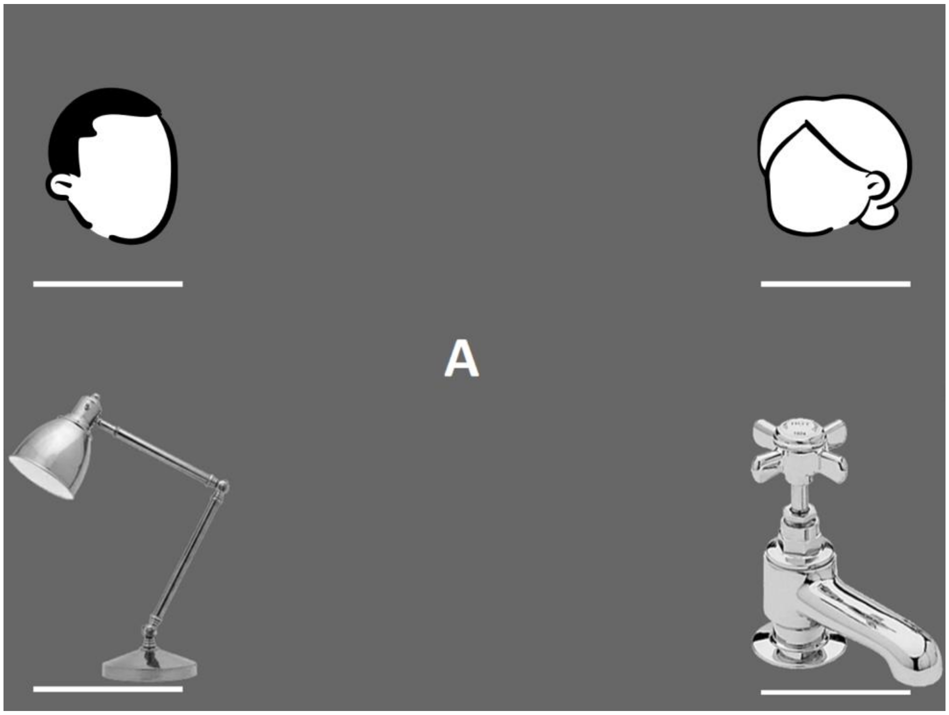


**Supplementary Fig. 1.** Illustration of the four images in the task: two faces and two objects, the actual faces are not shown in this illustration.

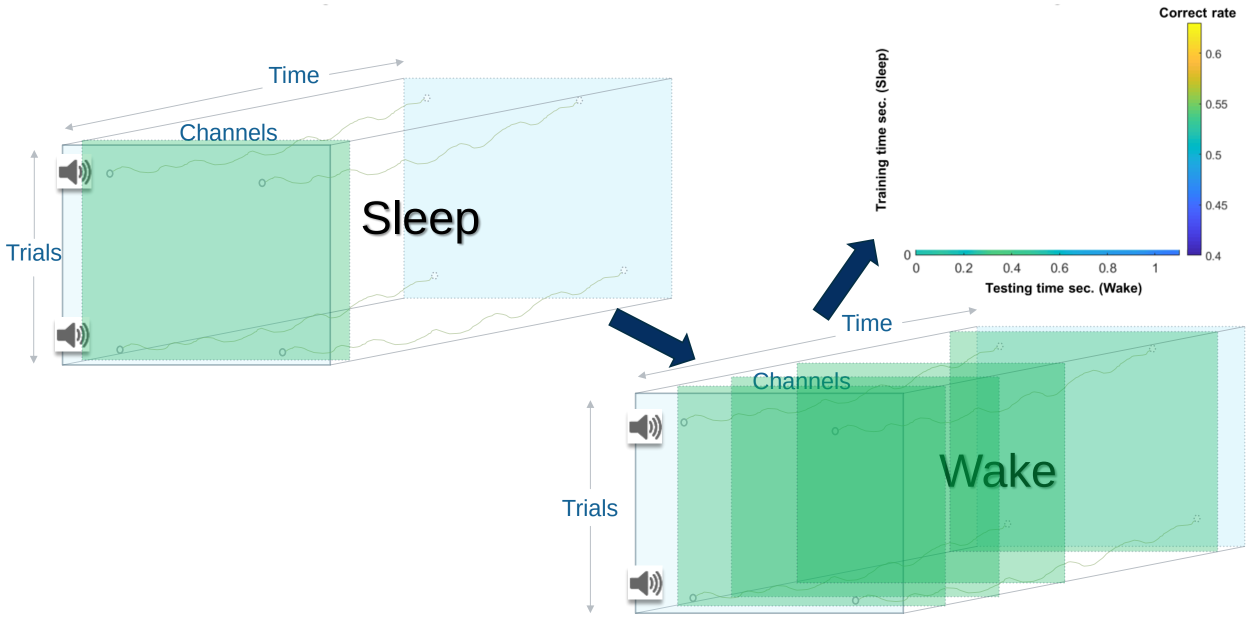


**Supplementary Fig. 2.** Example of the time x time classification procedure wherein one time point is used from sleep to build a classifier model and all wake time points were used for testing.

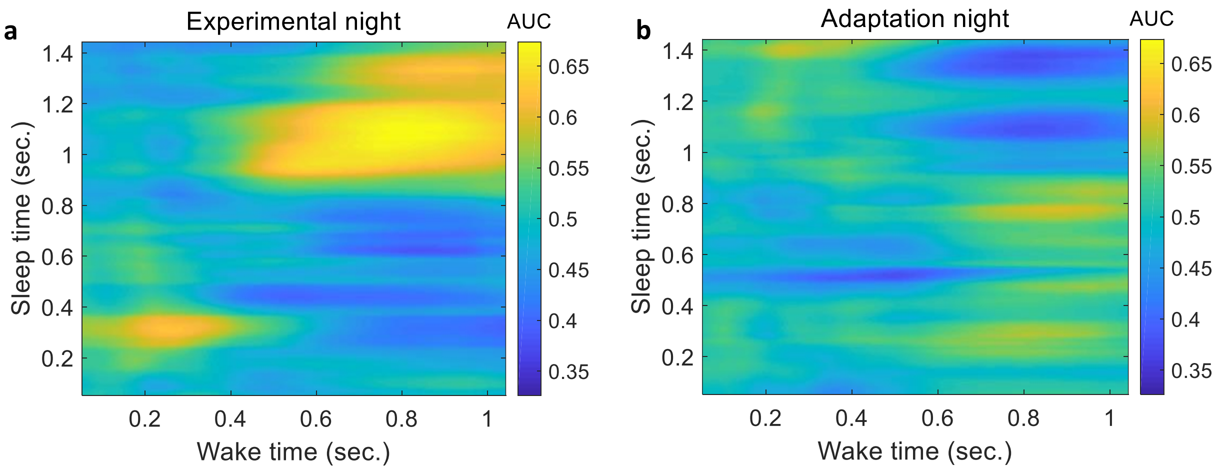

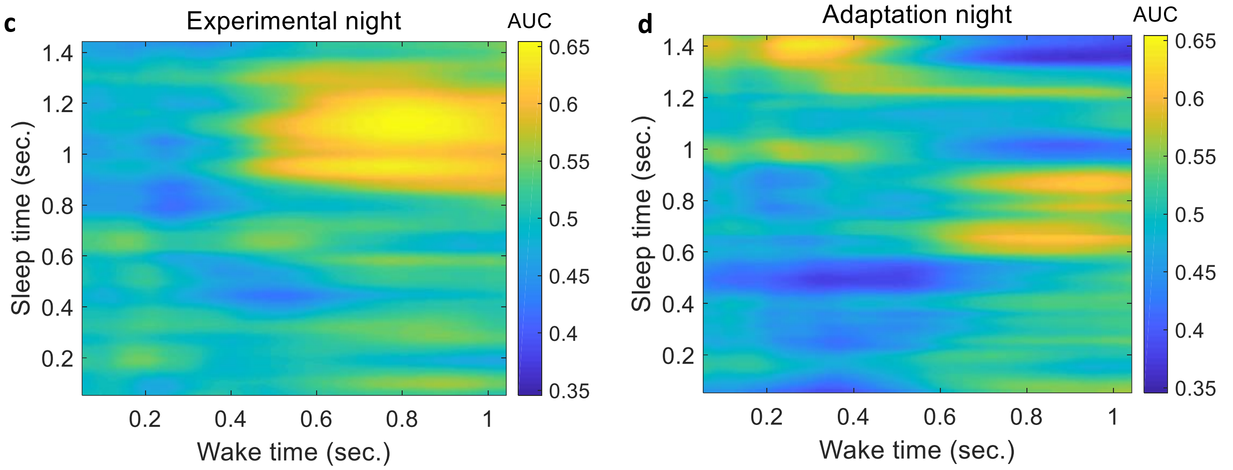


**Supplementary Fig. 3.** Classification of left- vs. right-hand. **a**, Classification performance when classifiers were trained on sleep using the experimental night and tested on wake. **b**, Classification performance when classifiers were trained on sleep using the adaptation night and tested on wake. **c**, Classification performance when classifiers were trained on sleep using the high theta power trials of the experimental night and tested on wake. **d**, Classification performance when classifiers were trained on sleep using the high theta power trials of the adaptation night and tested on wake.

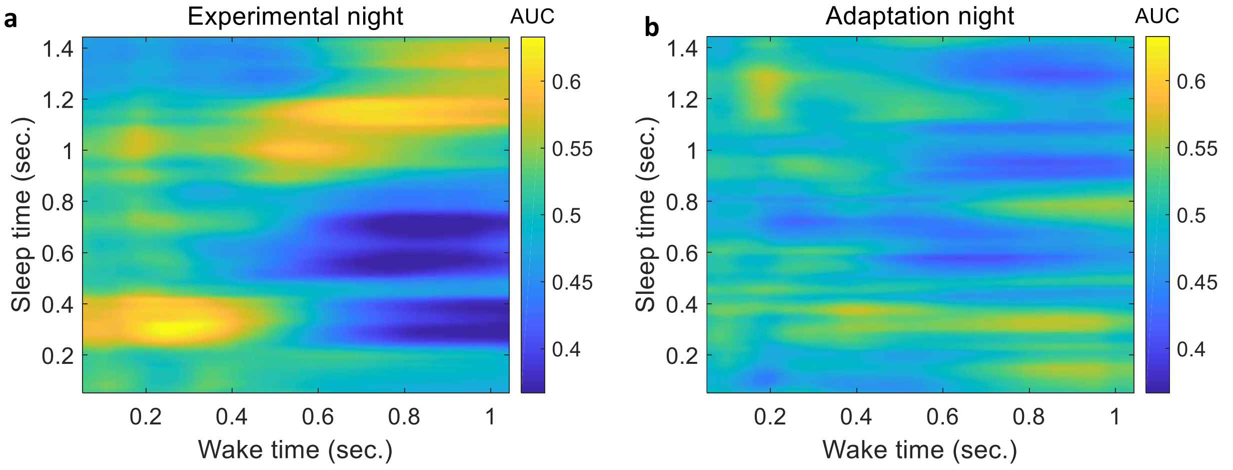

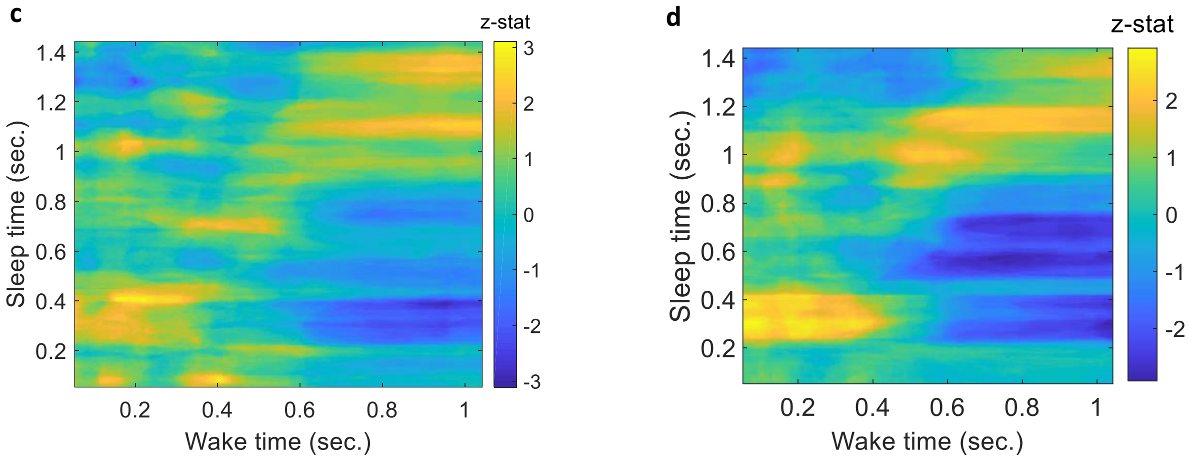


**Supplementary Fig. 4.** classification with low theta power trials. **a**, Classification of left- vs. right-hand using trials with low theta power using the experimental night did not show significant difference against the chance level, **b**, Likewise the classification using the adaptation night did not show significant difference against the chance level. **c**, Z-statistic values of the comparison between the classification of the experimental and the adaptation nights when the trials with low theta power were used. **d, c**, Z-statistic values of the comparison between the classification of the experimental and chance level.

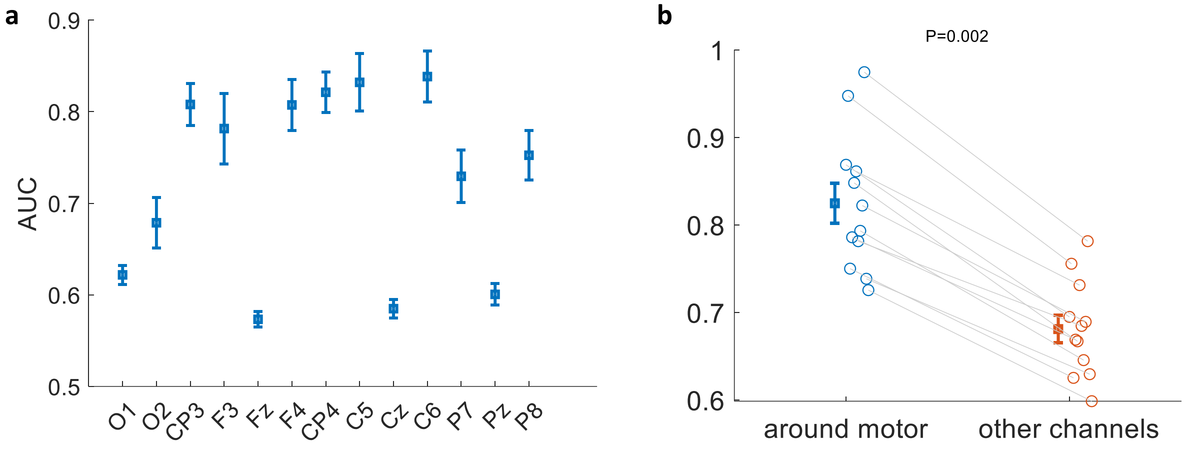


**Supplementary Fig. 5.** Searchlight analysis with linear classification on motor imagery data. **a**, Classification of left- vs. right-hand using searchlight analysis wherein timepoints from each channel are used as features to locate the channels with the highest classification performance, **b**, comparison of classification performance using the mean of the highest four channels (which came from channels around the motor area) and the mean of all other channels, which shows that the classification of those channels was significantly higher than the classification of all other channels.
